# Beyond Agreement: Standardizing Crowdsourced Synapse Annotations through Proofreading in EM Connectomics

**DOI:** 10.1101/2025.09.26.678851

**Authors:** Shi Yan Lee, Ana Correia, Nicolo Ceffa, Miranda Robbins, Daniel Franco-Barranco, Marta Zlatic, Albert Cardona, Samia Mohinta

## Abstract

Reliable synapse identification in volumetric EM is hampered by subtle, 3D cues that yield variable human judgments. We present a standardized proofreading protocol that pairs explicit, operational criteria with machine-learning candidate generation and a two-stage calibration of annotators. In two larval *Drosophila melanogaster* volumes imaged at 8×8×8 nm, five raters (expert + 4 calibrated annotators) reviewed model-proposed candidates using efficient node-based labels. Multi-rater judgments were aggregated with a probabilistic Dawid–Skene (DS) model to produce consensus labels with calibrated uncertainty. Post-calibration, individual annotator accuracy versus the expert improved (McNemar *p* < 0.05 for all raters), DS–expert agreement increased, and DS posterior entropy decreased for true positives/negatives, indicating more decisive consensus; gains were modest and dataset-dependent in chance-corrected agreement (Krippendorff’s *α*). By making uncertainty explicit, this protocol converts noisy judgments into auditable supervision suitable for training and evaluation, while honestly communicating residual ambiguity essential for reliable and robust connectomics at scale.

## 1 Introduction

Electron microscopy (EM) has rapidly advanced to generate terabyte–petabyte-scale [1, 2, 3, 4, 5] volumes at nanometer resolution, providing unprecedented access to the synaptic ultrastructure of neural tissue. These datasets are uniquely suited for reconstructing neural wiring diagrams (connectomes), a key step toward understanding how brain circuits give rise to behavior [6, 7, 8, 9, 10]. To meet the demands of such data scales, a range of automated machine-learning methods have emerged, enabling large-scale segmentation of neurons and detection of synapses directly from volumetric EM (vEM) data. Examples include U-Net variants [11, 12] and Flood-Filling Networks [13], which have substantially reduced the manual effort required for reconstruction. Despite these advances, automation remains far from self-sufficient: most methods depend on supervision from expert-annotated ground truth, which is both tedious to generate and exceedingly limited due to the scarcity of annotator time.

One response to the scarcity of expert annotations has been the adoption of crowdsourcing as a scalable alternative. By distributing labeling tasks to a large pool of non-expert contributors and aggregating their overlapping judgments, crowdsourcing can reduce the burden on individual annotators, mitigate the high costs of expert labor, and, when paired with appropriate quality control, approximate expert-level annotations at scale. This paradigm has gained traction in biomedicine and clinical sciences, powering applications from neuronal arbor reconstruction [14] and cell-morphology classification to lesion detection [15, 16], and increasingly supports AI pipelines that require large ground-truth datasets.

In connectomics, crowdsourcing has already demonstrated its value for scaling neuron reconstruction. EyeWire’s volunteers, for instance, contributed to Kim et al.’s seminal work on retinal motion-detection mechanisms [14], while the FlyWire consortium now organises a global community to proofread whole-brain reconstructions in fruit flies and other organisms, supported by infrastructure designed to coordinate many person-years of collective effort [17]. Web-based platforms such as CAVE [18] (with an adapted Neuroglancer viewer^1^), WebKnossos [19] and CATMAID [20] have been central to enabling such large-scale collaborative workflows. Yet agreement—even unanimous—does not ensure correctness; crowds can confidently converge on wrong answers [21], underscoring the need to model annotator reliability and to aggregate labels with principled methods rather than simple majority vote [22].

In practice, majority vote serves as a baseline, but reliability is typically audited with chance-corrected measures such as Fleiss’ *κ* and Krippendorff’s *α*; for complex, non-categorical annotations, the interpretability of *α* depends on the chosen distance function, motivating distributional/KS-based agreement metrics [23, 24]. Complementary variance-homogeneity checks (Levene’s; Brown–Forsythe ^2^) flag unstable per-annotator dispersion (e.g., time-per-item, confidence) before aggregation^3^ [25], and latent-truth estimation with Dawid–Skene leverages per-annotator confusion matrices to infer posterior labels, offering a principled upgrade over unweighted majority when expertise varies [26]. However, this consensus-and-reliability toolkit has seen limited evaluation for synaptic labels in vEM, where targets are small, ambiguous, and relational rather than purely categorical.

Here, we introduce a **standardized protocol** for synapse annotation through proofreading that yields reliable training targets for learning-based detectors. Specifically, we **(i)** define an operational criteria sheet for invertebrate (e.g., *Drosophila*) polyadic synapses to help annotators distinguish spurious from valid candidates; and **(ii)** adopt a consensus workflow that reports Krippendorff’s *α* to audit inter-annotator reliability and applies Dawid–Skene to aggregate annotator judgments into a probabilistically weighted consensus label for each candidate synapse. Crucially, our protocol makes uncertainty explicit – rather than obscured by raw agreement, and thereby reveals and quantifies the difficulty of achieving reliable consensus in manual synapse annotation through proofreading. Hence, we provide a transparent, reproducible pathway from multi-annotator judgments to training targets and establishes a practical proofreading protocol that others can adopt and extend.

## 2 Data and Task complexity

Synapse identification is difficult: synapses are sparse, irregular, boundary-poor, and their vEM appearance varies with staining, resolution, and image processing. Crucially, species and modality can change both morphology and topology, altering what annotators see and must decide.

### Vertebrates (anisotropic TEM)

Vertebrate synapses are reliably identified in transmission EM by canonical ultrastructural features, namely synaptic clefts, vesicle clusters, presynaptic active zones and postsynaptic densities [27]. These cues are well-resolved in large anisotropic datasets such as MICrONS [28, 29, 5], which trade z-resolution (typically ≤ 40 nm/px) for high in-plane sampling (typically ≤ 4 nm/px). Further aided by predominantly monadic (1-to-1) connectivity, synaptic partner assignment is straightforward in standard supervised pipelines.

### Invertebrates (isotropic FIB-SEM)

In invertebrate nervous systems, recent volumes are typically acquired with enhanced Focused Ion Beam–Scanning Electron Microscopy (eFIB-SEM [30]) at isotropic resolution [2, 31, 30] (e.g., 8 *×* 8 *×* 8 nm), vastly improving 3D continuity but reducing in-plane sharpness; as a result, clefts and post-synaptic densities (PSDs) can be faint (e.g., low-contrast) or noisy, complicating both manual labeling and training [27].

### Polyadic organization and its consequences

Insects frequently exhibit polyadic synapses in which a single presynaptic T-bar contacts multiple postsynaptic partners. In our data, this introduces topological ambiguity (how many post-synaptic partners?) and spatial ambiguity (which voxel neighborhood best captures the synaptic cleft and PSD constellation), both of which can be potential sources of inter-annotator disagreement and error propagation to learning systems.

### 3D verification

Synaptic features are inherently 3D. Both pre- and postsynaptic sites must be verified across multiple planes, since a postsynaptic density that is inconspicuous in one slice can become evident a few sections away. Practically, annotators verify synapses by scrolling along the *z*-axis in the *x*–*y* view and by consulting orthogonal reslices (*x*–*z, y*–*z*) to confirm the T-bar, cleft continuity, and the constellation of PSDs before assigning partners or rejecting the site. Ambiguity arises when features are indistinct (e.g., blurry or low-contrast), oblique to the imaging plane, or partially truncated; decisions in these cases often depend on an annotator’s prior experience and exposure to similar datasets.

Taken together, a reliance on tacit expertise, compounded by polyadic topology and appearance variability, is a major driver of inter-annotator disagreement when labeling synapses. These factors demand explicit, standardized criteria driving a consensus procedure to produce auditable ground truth for modeling and to systematically proofread machine-generated synapses before drawing biological conclusions. We demonstrate the successful application of our standardized protocol for synapse labeling by extracting, annotating, and analyzing multiple sub-volumes from two larval *Drosophila melanogaster* datasets acquired in-house using eFIB-SEM at 8*×*8*×*8 nm.

## 3 Materials and Methods

### 3.1 Synapse Labels and Baseline Detector

We used two whole-brain EM volumes from *Drosophila melanogaster* larvae—**Octo** (first instar, L1) and **MR** (second instar, L2), prepared by chemical fixation and imaged with eFIB-SEM at isotropic 8×8×8 nm resolution. The datasets were registered and post-processed with CLAHE (parameters: block radius 200, 256 bins, slope 2.0), and served in CATMAID for manual synapse labeling/proofreading.

A domain expert (∼ 20 years’ experience) produced the seed gold-standard labels (ground-truth) only on Octo, following the conventions introduced in this work (Table 1), i.e., a presynaptic node on the presynaptic neurite, a connector on the T-bar, and postsynaptic nodes on each target neurite at the visible PSD. To concentrate effort and mitigate sparsity, the expert annotated three 5 *×* 5 *×* 5 µm volumes (∼ 625^3^-voxel cubes at 8nm resolution).

**Table 1:**
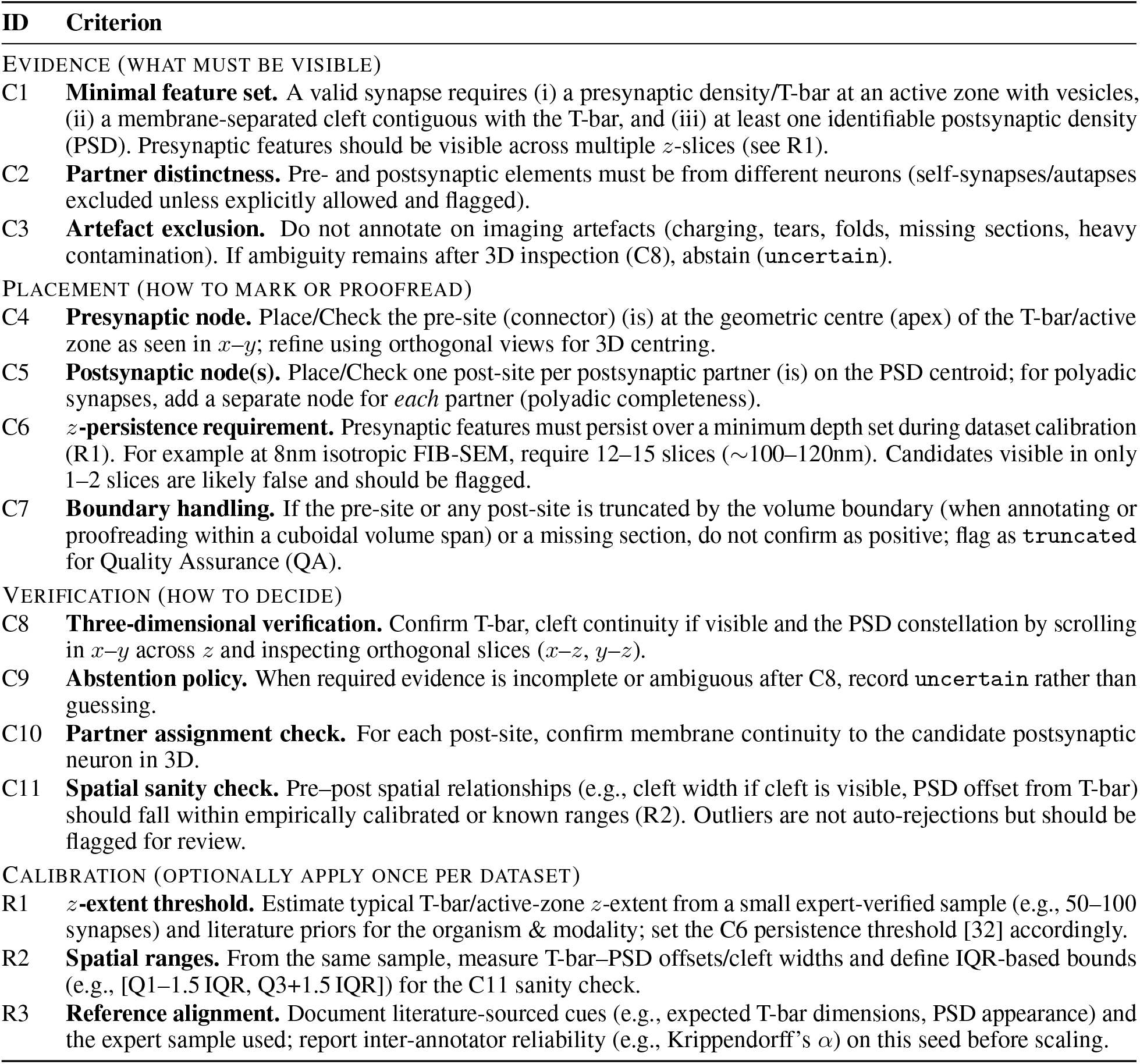
Standardized protocol for synapse annotation and proofreading (node-based annotations; applicable to CATMAID/CAVE-style tools).

| ID | Criterion |
| --- | --- |
| EVIDENCE (WHAT MUST BE VISIBLE) |  |
| C1 | **Minimal feature set.** A valid synapse requires (i) a presynaptic density/T-bar at an active zone with vesicles, (ii) a membrane-separated cleft contiguous with the T-bar, and (iii) at least one identifiable postsynaptic density (PSD). Presynaptic features should be visible across multiple $z$-slices (see R1). |
| C2 | **Partner distinctness.** Pre- and postsynaptic elements must be from different neurons (self-synapses/autapses excluded unless explicitly allowed and flagged). |
| C3 | **Artefact exclusion.** Do not annotate on imaging artefacts (charging, tears, folds, missing sections, heavy contamination). If ambiguity remains after 3D inspection (C8), abstain (uncertain). |
| PLACEMENT (HOW TO MARK OR PROOFREAD) |  |
| C4 | **Presynaptic node.** Place/Check the pre-site (connector) (is) at the geometric centre (apex) of the T-bar/active zone as seen in $x$-$y$; refine using orthogonal views for 3D centring. |
| C5 | **Postsynaptic node(s).** Place/Check one post-site per postsynaptic partner (is) on the PSD centroid; for polyadic synapses, add a separate node for *each* partner (polyadic completeness). |
| C6 | **$z$-persistence requirement.** Presynaptic features must persist over a minimum depth set during dataset calibration (R1). For example at 8nm isotropic FIB-SEM, require 12–15 slices ($\sim 100$–$120$nm). Candidates visible in only 1–2 slices are likely false and should be flagged. |
| C7 | **Boundary handling.** If the pre-site or any post-site is truncated by the volume boundary (when annotating or proofreading within a cuboidal volume span) or a missing section, do not confirm as positive; flag as truncated for Quality Assurance (QA). |
| VERIFICATION (HOW TO DECIDE) |  |
| C8 | **Three-dimensional verification.** Confirm T-bar, cleft continuity if visible and the PSD constellation by scrolling in $x$-$y$ across $z$ and inspecting orthogonal slices ($x$-$z$, $y$-$z$). |
| C9 | **Abstention policy.** When required evidence is incomplete or ambiguous after C8, record uncertain rather than guessing. |
| C10 | **Partner assignment check.** For each post-site, confirm membrane continuity to the candidate postsynaptic neuron in 3D. |
| C11 | **Spatial sanity check.** Pre–post spatial relationships (e.g., cleft width if cleft is visible, PSD offset from T-bar) should fall within empirically calibrated or known ranges (R2). Outliers are not auto-rejections but should be flagged for review. |
| CALIBRATION (OPTIONALLY APPLY ONCE PER DATASET) |  |
| R1 | **$z$-extent threshold.** Estimate typical T-bar/active-zone $z$-extent from a small expert-verified sample (e.g., 50–100 synapses) and literature priors for the organism & modality; set the C6 persistence threshold [32] accordingly. |
| R2 | **Spatial ranges.** From the same sample, measure T-bar–PSD offsets/cleft widths and define IQR-based bounds (e.g., [Q1–1.5 IQR, Q3+1.5 IQR]) for the C11 sanity check. |
| R3 | **Reference alignment.** Document literature-sourced cues (e.g., expected T-bar dimensions, PSD appearance) and the expert sample used; report inter-annotator reliability (e.g., Krippendorff’s $\alpha$) on this seed before scaling. |

Using these annotated Octo sub-volumes, we trained a preliminary 3D presynaptic-site detector [33, 34] with connector (T-bar) coordinates as presynaptic targets and the post-synaptic node coordinates are post-synaptic targets. We then re-ran inference on the same Octo cubes to surface potential missed synapses. Importantly, non-matching predictions were treated as candidate false positives (FPs) only under the detector’s matching rule (for F1 score calculations [34]); many such candidates are plausible true synapses that were (i) excluded by protocol (e.g., boundary-truncated sites not permitted by Table 1), or (ii) missed in the seed owing to appearance variability and task complexity.

The model was not trained on MR; instead, we applied the Octo-trained detector directly to MR to propose synapse candidates (OOD evaluation). We earmarked two 4 *×* 4 *×* 4 µm (500^3^ -voxel) MR sub-volumes for this purpose and, for practicality, ran inference on one cube; all predictions from that cube were triaged for proofreading to rapidly generate MR ground truth. For the multi-rater study, we reviewed predictions from one of the three Octo cubes and one of the two MR cubes (i.e., 1/3 Octo and 1/2 MR), reflecting the hundreds of synapses per cube and limited annotator availability.

All candidate FPs (Octo and MR) were imported into cloned CATMAID instances with the original Octo ground-truth withheld to avoid anchoring bias. Five raters—four biologists (6 months–5 years’ experience) and the expert (∼ 20 years)—independently reviewed the same candidates and initially assigned one of three labels: correct, uncertain, or incorrect. These judgments were subsequently used to (i) quantify inter-annotator reliability (e.g., majority vote, Krippendorff’s *α*) and (ii) aggregate labels into a probabilistic consensus via Dawid–Skene.

### 3.2 Consensus models and reliability statistics

We quantify consensus across five raters: the expert annotator (`gs`; *gold standard*) and four additional annotators with varied EM synapse experience (6 months–5 years). Synapse candidates are labeled as {`correct, incorrect, uncertain`}. Below we describe the aggregation and agreement measures used.

#### Majority voting (baseline)

As a baseline aggregation method, we use simple majority voting (`mv`). For each item *j* with annotations 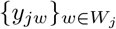 and class set *C* = {`correct, incorrect, uncertain`}, the aggregated label is the most frequent class:

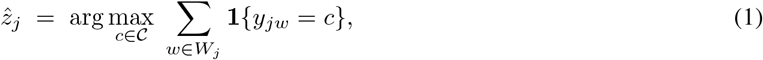

where **1** {·} is the indicator function. In case of a tie (two or more classes with equal maximum votes), we assign the label `uncertain`.

#### Dawid–Skene (DS) model

We aggregate annotator judgments with the Dawid–Skene model [26], which posits a latent true class *z*_*j*_ ∈ *C* for each item *j* and annotator–specific noise via confusion matrices. Let *C* = {1, …, *K*} denote the class set (binary: {`correct, incorrect`}; tri-class: {`correct, uncertain, incorrect`}), and let *W*_*j*_ be the annotators who labeled item *j*. Annotator *w* supplies an observed label *y*_*jw*_ ∈ *C*. Each annotator *w* has a *K × K* confusion matrix *e*^(*w*)^ with entries 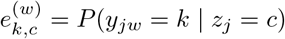 and 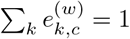 for each *c*. Class priors are *π*_*c*_ = *P* (*z*_*j*_ = *c*) with ∑_*c*_ *π*_*c*_ = 1. Conditional on *z*_*j*_, rater responses are assumed independent.

The model is fit via the EM algorithm [26], yielding item-wise posteriors

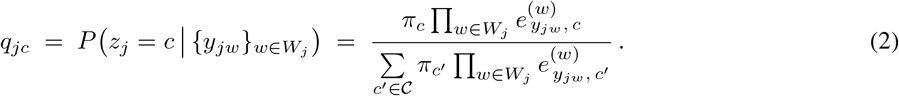

In this expression, 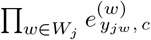 is the likelihood of the observed annotations for item *j* assuming true class *c*, and the denominator normalizes over all *c*^*′*^ ∈ *C* so that ∑_*c*_ *q*_*jc*_ = 1.

The posterior vector *q*_*j*_ = (*q*_*jc*_)_*c*∈*C*_ can be either used as a *soft* consensus or as a *hard* consensus:

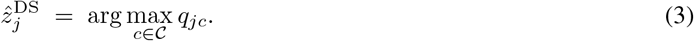

#### Skill-weighted Dawid–Skene (SW-DS)

To account for differences in annotator performance, we introduce fixed *skill weights s*_*w*_ ∈ [0, 1] for each annotator *w*, representing reliability estimated relative to a reference set (gold standard, gs). In DS, the weights modulate each annotator’s likelihood contribution via exponentiation:

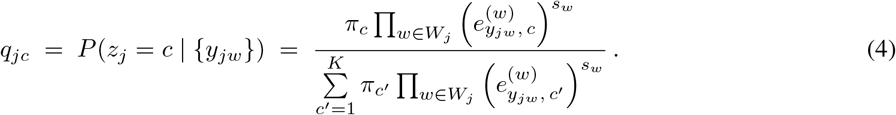

High-skill annotators (*s*_*w*_ ≈ 1) exert greater influence on the aggregated label distribution, whereas lower-skill annotators (*s*_*w*_ ≈ 0) contribute proportionally less.

#### Entropy (uncertainty)

For each item *j*, DS produces a posterior *q*_*j*_ = (*q*_*j*1_, …, *q*_*jK*_), with *q*_*jc*_ = *P* (*z*_*j*_ = *c* | {*y*_*jw*_}). We quantify aggregation uncertainty using Shannon entropy:

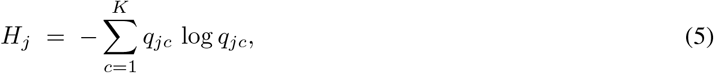

Higher *H*_*j*_ indicates greater uncertainty (posterior mass spread across classes).

#### Krippendorff’s *α* (interval)

We map labels to numeric scores *x* ∈ {0, 0.5, 1} via `incorrect` ⟼ 0, `uncertain` ⟼ 0.5, and `correct` ⟼ 1, and use the squared distance *δ*^2^(*x, y*) = (*x − y*)^2^. Let items (units) be *u* = 1, …, *U*, with *x*_*uk*_ the score given by annotator *k* to unit *u, n*_*u*_ the number of annotations (synapse classifications) for unit *u*, and *N* = ∑_*u*_ *n*_*u*_ the total number of annotations. Observed disagreement averages within-unit pairwise squared differences:

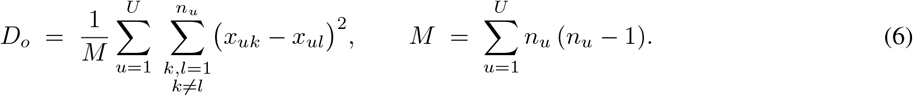

Expected disagreement averages pairwise squared differences across all annotations (chance level):

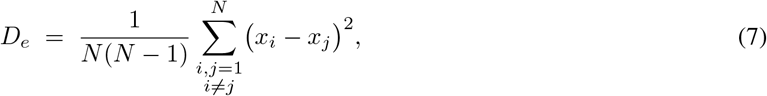

where *x*_*i*_ indexes the pooled list of all annotations. Krippendorff’s alpha (interval) is

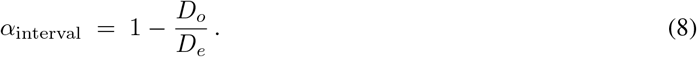

We report *α* overall and for subsets filtered by the expert’s `gs` label (*gs* ∈ {`correct, incorrect, uncertain`}). Empty subsets (no pairable annotations) yield undefined *α*.

#### Pre/post reliability change

To assess whether proofreading improved annotation reliability, we compute

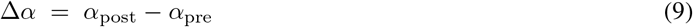

Uncertainty is quantified via nonparametric bootstrapping: items are resampled with replacement (*B* = 2000 replicates), Δ*α* is recomputed per replicate, and we report the mean difference, a 95% percentile confidence interval, and a one-sided *p*-value (*p* = Pr[Δ*α ≥* 0]).

#### McNemar’s significance test

For paired categorical comparisons (e.g., pre/post accuracy or aggregated DS labels), we apply McNemar’s test on discordant pairs to assess whether changes are systematic. Results are visualized with right-side significance brackets; asterisks indicate significance levels as defined in the figure legends.

### 3.3 Crowdsourced proofreading workflow

We divided the review process into two stages with a calibration in between. Five raters participated: the expert annotator (gs; *gold-standard*) and four additional annotators with varied experience in EM synapse labeling (6 months–5 years). Henceforth, we refer to the four non-expert raters as `calibrated annotators` (CAs; CA1, CA2, CA3 and CA4) to reflect that they received protocol training and took part in a calibration step before formal review. Fig. 1 illustrates the proofreading instructions followed by CAs and the expert annotator.

**Figure 1:**
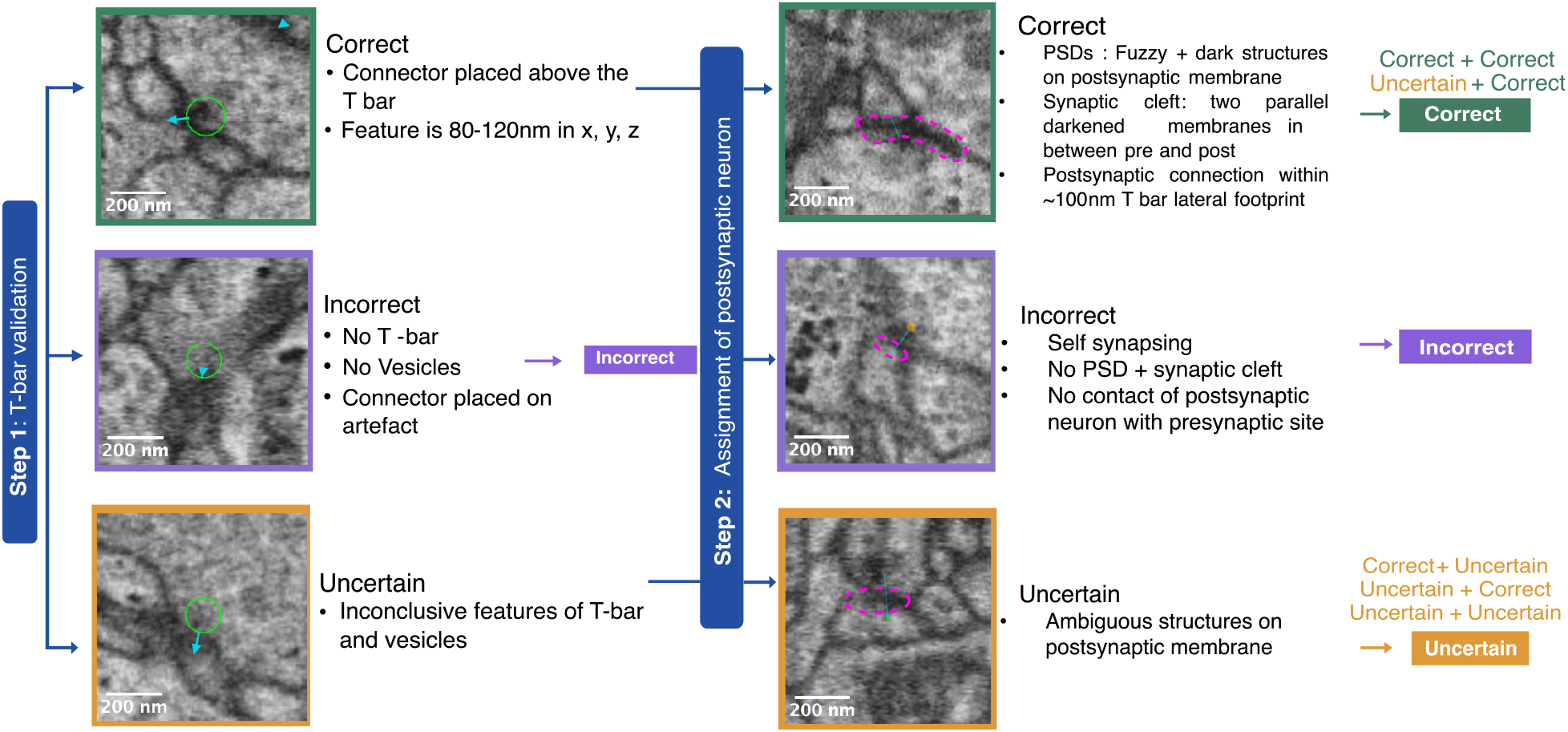
Initial proofreading instructions on synapse classification: The presence of a T-bar within the connector region (green circle) at the active zone was first evaluated, and synapses were classified into one of three categories to guide the next step. Only those marked as `correct` or `uncertain` proceeded to the subsequent stage of evaluation. In these cases, the postsynaptic neuron was assessed, represented by the traced magenta area. The final classification was determined by combining presynaptic and postsynaptic assignments, with all possible outcomes summarized in the far-right column. Each synapse was ultimately categorized as `correct, incorrect`, or `uncertain`.

#### Stage 1 — initial calibration and first-pass review

After a short training session involving the CAs that introduced the protocol, examples and edge cases (Fig. 1), **all** candidate synapses from **Octo** and **MR** were distributed for independent review. Each annotator independently classified every candidate as `correct, incorrect`, or `uncertain`, working asynchronously at their own pace in cloned CATMAID instances (with gold-standard labels withheld).

*Consensus modeling*. From per-synapse, multi-rater judgments we derived consensus labels using:

1. Majority vote over the three classes.
2. Dawid–Skene (DS) latent-truth estimation to learn per-annotator confusion matrices and infer posterior probabilities for `correct, uncertain, incorrect`; we instantiated DS on different rater pools and a skill-weighted variant (see section 3.2) that uses estimated annotator reliabilities as priors.
3. Thresholding and uncertainty summaries: items were assigned to a class by arg max posterior, with posterior entropy retained as an item-level uncertainty metric. For auditing and rater feedback we computed: (i) normalized per-annotator label distributions (class propensity heatmap); (ii) per-annotator confusion matrices against a reference (expert or DS consensus; see supplementary Fig. S1); and (iii) an agreement-vs-experience analysis using a simple linear fit with 95% confidence band (experience recorded in years).

#### Stage 2 — second calibration session and post-calibration evaluation

Following consensus derivation, we held a second training session with the expert to walk through borderline cases and reaffirm the criteria in Table 1. We then ran a post-consensus evaluation under a simplified binary task (labels restricted to `correct` or `incorrect`) to test whether probabilistic consensus improves reliability when the uncertain option is removed. For this post-test, we sampled from both cubes. From the MR cube, a balanced set of 90 synapses (45 previously classified as `correct`, 45 as `incorrect` by `gs`). From the Octo cube, a similar set was initially selected; however, some candidates were subsequently discarded, leaving a final set of 46 synapses. In total, 136 synapses were redistributed to all annotators. We compared Stage 1 and post-calibration (Stage 2) outcomes using paired analyses, including McNemar’s test on the binary outcomes to assess shifts in error patterns, and recomputed DS posterior entropy to summarize uncertainty on the binary task.

## 4 Results

### 4.1 Consensus after Stage 1 Calibration

Following the initial calibration phase, we first sought to characterize the behavior of the individual annotators and establish a baseline for consensus. Fig. 2 presents a comprehensive analysis of rater agreement and the impact of different aggregation methods on the initial set of **Octo** and **MR** candidates. This analysis reveals significant variability among raters and demonstrates how modeling choices can systematically shift the final consensus labels.

**Figure 2:**
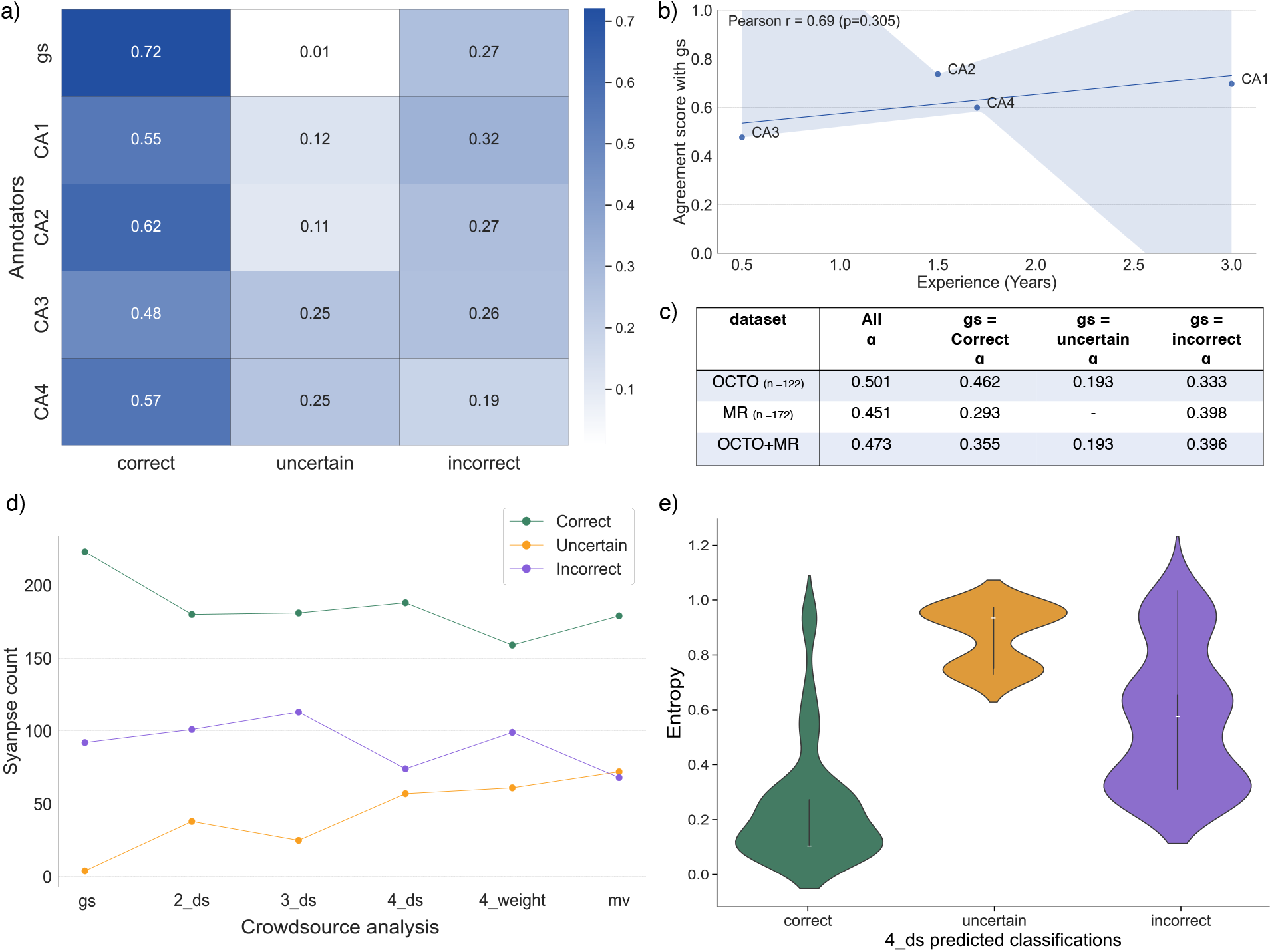
Annotator behavior and consensus after Stage 1 calibration. **a)** Row-normalized heatmap showing per-rater label usage, highlighting different decision thresholds. **b)** Per-rater agreement score plotted against years of experience, showing a modest positive correlation. **c)** Krippendorff’s *α* reliability, calculated overall and stratified by dataset and the expert’s (gs) label. **d)** Effect of aggregation method on the final count of `correct, uncertain`, and `incorrect` labels. Legend note: `gs` = expert-only reference (gold standard); `k_ds` = unweighted DS with *k* raters and `4_weight` = skill-weighted DS. Majority vote baseline is denoted as mv. **e)** Distribution of model uncertainty (posterior entropy) for each consensus label (lower is better).

#### Annotator propensities

Fig. 2a reveals substantial differences in label usage across raters, indicating systematic differences in caution/thresholds. The expert (`gs`) rarely selects `uncertain` (∼0.01) and favors `correct` (∼0.72), with the remainder `incorrect` (∼0.27). By contrast, `CA3` and `CA4` use `uncertain` far more often (∼0.25), with `correct` spanning ∼0.48–0.57 and `incorrect` ranging from ∼0.26 (`CA3`) to ∼0.19 (`CA4`). The remaining raters (`CA1, CA2`) fall between these extremes. Because the heatmap is row-normalized, these differences reflect per-rater propensities rather than class prevalence in the data.

#### Agreement vs experience

Fig. 2b relates the CAs’ years of EM/synapse experience to their agreement score with gs labels. The slope is shallowly positive, indicating that greater experience is associated with slightly higher agreement with gs, but the effect size is modest, consistent with the task remaining difficult even for trained raters.

#### Agreement by dataset and class

Fig. 2c reports Krippendorff’s *α* per dataset (e.g., Octo, MR) and pooled, alongside per-class *α*. Overall *α* falls in the low–moderate range [35] and varies by dataset. Per-class *α* is lower and more variable, especially for `correct/incorrect`, reflecting that much of the disagreement sits on the boundary between `correct` and `incorrect` (with many raters opting for `uncertain`), and that `uncertain` agreements tend to be weak.

#### Effect of aggregation setting on class balance

Fig. 2d illustrates how the counts/proportions of items assigned to `correct, uncertain`, and `incorrect` change across aggregation settings (`gs, 2_ds, 3_ds, 4_ds, 4_weight`). Moving from gs toward Dawid–Skene with more raters and skill-weighted DS (see section 3.2), the share of `uncertain` increases, `correct` decreases modestly, and `incorrect` changes less. This pattern reflects the conservative effect of applying skill weights in DS, which down-weights less reliable raters and makes ambiguity explicit (see Supplementary Fig. S2).

#### Posterior uncertainty by class

Fig. 2e shows Dawid–Skene (un-weighted) posterior entropy by final label. Entropy is lowest for `correct`, higher for `uncertain`, and highest/broadest for `incorrect`, consistent with more decisive posteriors for items judged `correct` and more diffuse posteriors for items judged `incorrect`.

In summary, the analysis of Stage 1 confirmed that the proofreading task was intrinsically difficult, characterized by substantial differences in rater decision thresholds. Merely identifying this ambiguity through statistical aggregation was insufficient for our goals. Therefore, these findings established the necessity for a second calibration stage focused on active reconciliation, aiming to move beyond statistical modeling to improve rater alignment and the definitiveness of the final labels.

### 4.2 Consensus after Stage 2 Calibration

To achieve the goal of active reconciliation and improve rater alignment, we implemented the second calibration stage. This intervention involved a focused re-review of 136 synapses (46 from the Octo dataset and 90 from MR), where annotators applied the reinforced proofreading criteria under a simplified binary classification task. Fig. 3 evaluates the success of this approach, illustrating the resulting changes in annotator behavior and consensus reliability.

**Figure 3:**
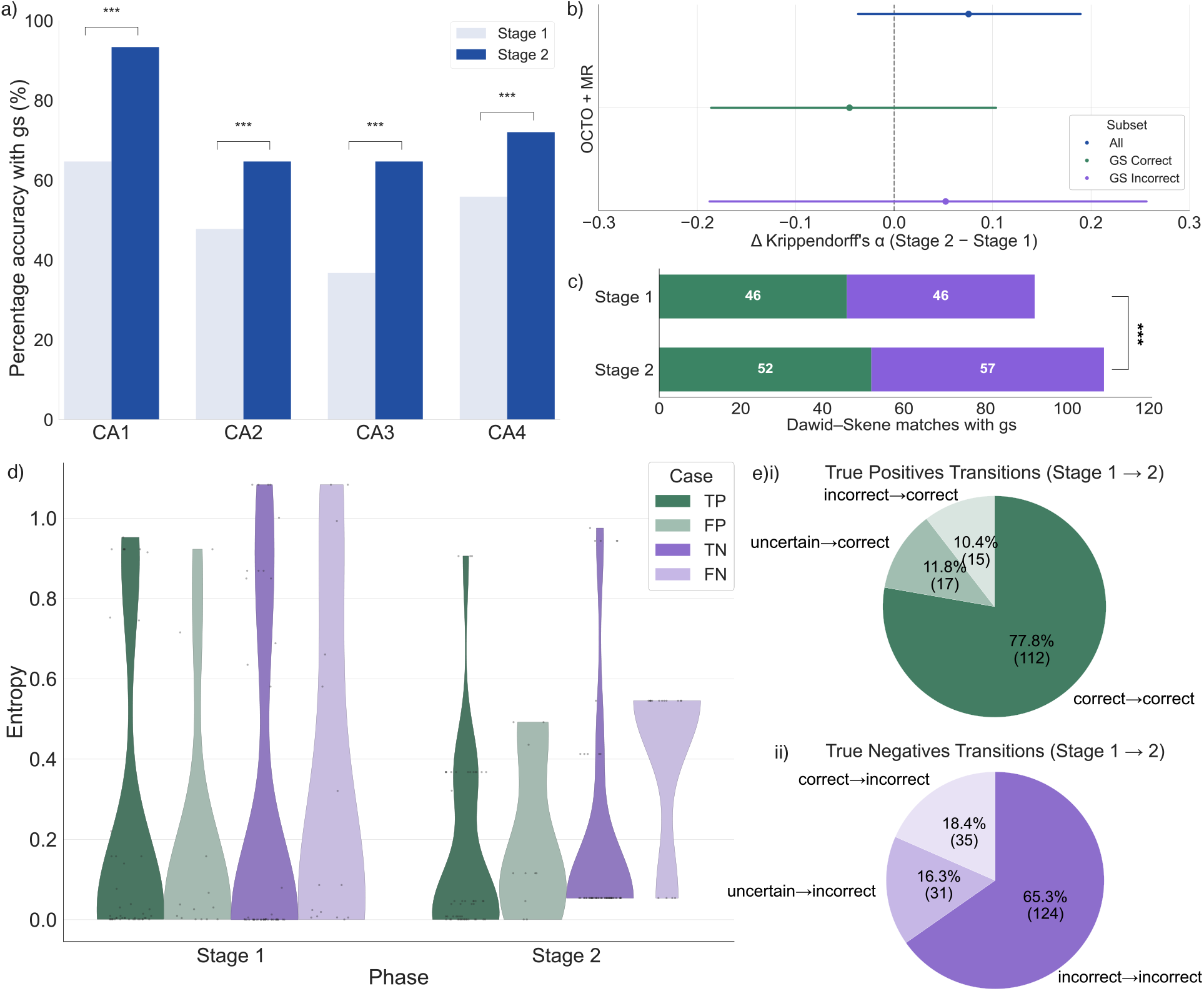
Annotator Alignment and Consensus Reliability After Stage 2 Calibration. **a)** Significant improvement in accuracy against the gold standard (gs) for all annotators. **b)** Change in Krippendorff’s *α*, showing a positive consensus shift overall and particularly for synapses previously labeled `incorrect` by the expert (gs). **c)** Dawid–Skene consensus labels from calibrated annotators (4) show significantly improved alignment with the gold standard (1 expert) after calibration. **d)** Dawid–Skene posterior entropy decreases for concordant cases (True Positives (TP) and True Negatives (TN)), indicating more decisive and confident consensus. **e)** Transition of labels for items ultimately classified as (i) True Positives and (ii) True Negatives, showing that the calibration helped resolve many previously `uncertain` cases.

#### Rater-level agreement and pairwise alignment

Under the binary post-test, overall agreement with the gold standard (gs) improved for every annotator (McNemar’s test on paired outcomes, *p <* 0.05 for all; Fig. 3a). The largest gains were observed for `CA1` and `CA3`, consistent with a shift toward gs’s decision criteria after the second calibration. Concordantly, F1 scores of `CA1` and `CA3` versus gs increased(Supplementary Fig. S4), indicating improved precision–recall trade-offs rather than a mere change in class balance for those individuals. Pairwise alignment gains were heterogeneous across annotators, with the largest improvements concentrated in a subset of raters (Supplementary Fig. S3).

#### Group-level agreements and consensus structure

Chance-`correct`ed agreement, quantified by Krippendorff’s *α* showed no significant overall shift after Stage 2, with only small positive changes for the full set (Δ ≈ 0.076) and for items labeled `incorrect` by gs (Δ ≈ 0.052) (Fig. 3b, also see Supplementary Fig. S5).

Applying Dawid–Skene sharpened consensus: post-calibration, the DS assignment matched gs more often overall, with the largest gains in the `incorrect` class (Fig. 3c). This aligns with the individual-level patterns (Fig. 3a), indicating that recalibration primarily `correct`ed previously error-prone decisions.

Posterior `uncertain`ty from DS decreased where it mattered: entropy was lower for true positives and true negatives after Stage 2, indicating more decisive consensus (Fig. 3d, Supplementary Fig. S6). By contrast, false positives and false negatives concentrated at intermediate entropies (≈ 0.60), suggesting that raters were often confident yet wrong on a residual set of difficult cases.

#### Effect of removing the uncertain option

Before proofreading, elevated use of `uncertain` made definitive assignment difficult. In the post-proofreading (binary) adjudication, we tracked how previously `uncertain` items resolved relative to the gold standard (gs). Among these, 11.8% converted to true positives (Fig. 3e(i)) and 18.3% to true negatives (Fig. 3e(ii)), indicating that binary adjudication helped resolve a substantial fraction of ambiguous cases in the `correct` direction. Fig. 3e summarizes all label transitions (including `incorrect` ⟼ `correct` and `correct` ⟼ `incorrect`), highlighting the magnitude of judgment shifts that collectively underlie the observed post-proofreading accuracy gains (also see Supplementary Fig. S8).

## 5 Discussion

We provide, to our knowledge, one of the first systematic characterizations of inter-annotator variability in EM synapse annotation through proofreading together with a standardized, consensus-centered pipeline that makes reliability auditable and `uncertain`ty explicit.

### Annotator subjectivity persists, even with written guidance

Even with a visual guideline and an initial calibration, raters exhibited distinct decision thresholds, evident in heterogeneous use of the `uncertain` label and modest, experience-related gains. Cases that the expert (gs) considered straightforward were frequently labeled `uncertain` by less-experienced raters, increasing ambiguity at the pool level. This reflects the task complexity and indicates that written guidelines, by themselves, may not remove subjective decision boundaries.

### Structured recalibration improved alignment but not uniformly

Stage 2 recalibration increased per-annotator agreement with gs across the board and raised F1 scores (Supplementary Fig. S4), with the largest gains for raters who were most conservative at Stage 1. Pairwise alignment also improved for most rater pairs (see Supplementary Fig. S3), though one rater remained comparatively idiosyncratic. Removing the `uncertain` option in the post-test resolved a meaningful fraction of previously ambiguous items in the `correct` direction (into gs-consistent true positives or true negatives), suggesting that forced decisions, after targeted calibration, can reduce residual indecision without wholesale degradation of accuracy.

### Probabilistic aggregation yields a more robust consensus and calibrated uncertainty

Dawid–Skene (DS) consensus increased agreement with `gs`. Stratified by class, the largest gains occurred in the `incorrect` subset, suggesting improved sensitivity to erroneous synaptic features. However, a supplementary analysis of rationale codes (see Supplementary Fig. S7) suggests that even when consensus improves, annotator reasons for `incorrect` calls are not always aligned. Across stages, DS entropy behaved consistently: in Stage 1 it was lowest for `correct` and highest/broadest for `incorrect`; after recalibration in Stage 2 it decreased for true positives and true negatives, whereas false positives/negatives concentrated at intermediate values, separating certainty profiles between decisive and contentious cases. In our data, unweighted DS outperformed majority vote and a simple skill-weighted variant on the same items, yielding more gs-consistent assignments while retaining useful `uncertain`ty summaries; majority vote ignores rater reliability, and naïve weighting can overemphasize individual biases on difficult items. (Note: gs labels were used only for evaluation, not to train DS.)

### Practical limits: ambiguity remains

Chance-`correct`ed agreement (Krippendorff’s *α*) remained low–moderate overall and was dataset-dependent; gains were small and uneven across strata. Note that post Stage 2 evaluation Octo analyses were underpowered relative to MR (46 vs. 90 items), which likely increased variance and masked consensus gains. False-positive and false-negative subsets retained intermediate entropies, indicating confident but wrong judgments on a residual set of borderline synapses. These observations bound what consensus can achieve: when ultrastructural evidence is intrinsically weak or truncated, no amount of pooling fully removes ambiguity. Furthermore, demonstrating high reliability is challenging with a conservative metric like Krippendorff’s alpha, which penalizes any deviation from perfect consensus and is strongly depressed by even a single dissenting score.

Although this study was limited to two larval *Drosophila* sub-volumes and a small rater cohort, we show that calibrated guidance combined with probabilistic consensus provides an auditable path from noisy human judgments to reliable supervision. This protocol offers a principled way to manage and communicate residual `uncertain`ty, an essential step for advancing the reliability of connectomics.

## 6 Conclusion and Future Directions

Crowdsourced proofreading of synapses can yield auditable, consensus-grade supervision for synapse detection, yet subjectivity and ambiguity persist – most acutely on borderline cases. By pairing calibrated guidance with probabilistic consensus (Dawid–Skene with `uncertain`ty retained), our protocol transparently converts multi-annotator judgments into training targets while keeping `uncertain`ty explicit, enabling safer model training and more trustworthy biological readouts. Future work will explore extensions in scale and diversity (more raters and volumes), generalization across organisms and modalities, and mitigation of selection effects via unbiased sampling. In the near term, we advocate using consensus/model `uncertain`ty (e.g., DS entropy) to prioritize reviews, stratifying tasks by difficulty to focus calibration, and training detectors on soft consensus labels and disagreement structure rather than hard labels alone. In short, our protocol confronts the core challenge—subjective, noisy judgments—by turning it into calibrated, `uncertain`ty-explicit supervision, supporting reliable and reproducible connectomics as datasets and teams scale.

## 7 Acknowledgments

We are grateful to Marc Corrales and Nadine Randel (MRC LMB), and to C. Shan Xu and Song Pang (HHMI Janelia Research Campus, Yale University), for acquiring the MR and Octo volumes. We also thank Michael Clayton (MRC LMB), Amina Dulac and Bernd Breuer (University of Cambridge) for valuable discussions.

## 8 Funding

This project was funded by a Wellcome Trust Investigator Award to Albert Cardona (Ref: 205038/Z/16/Z), an ERC Grant to Marta Zlatic (Ref: ERC-2018-COG: 819650), a BBSRC Grant to Albert Cardona (Ref: APP26929, Population Connectomics) and the MRC LMB core funding.

## 9 Author contributions

The author contributions are as follows:

Conceptualization: SYL and SM; Methodology: SYL, AC^b^, NC, MR; Investigation: SYL; Visualization: SYL, DFB, SM; Supervision: SM, AC^a^; Writing—original draft: SYL and SM; Writing—review and editing: SYL, SM, DFB, AC^b^, MR and AC^a^. Note: AC^b^: Ana Correia, AC^a^: Albert Cardona

## 10 Competing Interests

The authors declare no competing interests.

## Footnotes

1 https://github.com/google/neuroglancer.git

2 https://influentialpoints.com/Training/levenes_test.htm

3 https://www.itl.nist.gov/div898/handbook/eda/section3/eda35a.htm

## Notes

### Competing Interest Statement

The authors have declared no competing interest.

### Summary of Updates

This version contains updated funding information and acknowledgements.

