## Supplementary Material for "Beyond Agreement: Standardizing Crowdsourced Synapse Annotations through Proofreading in EM Connectomics"

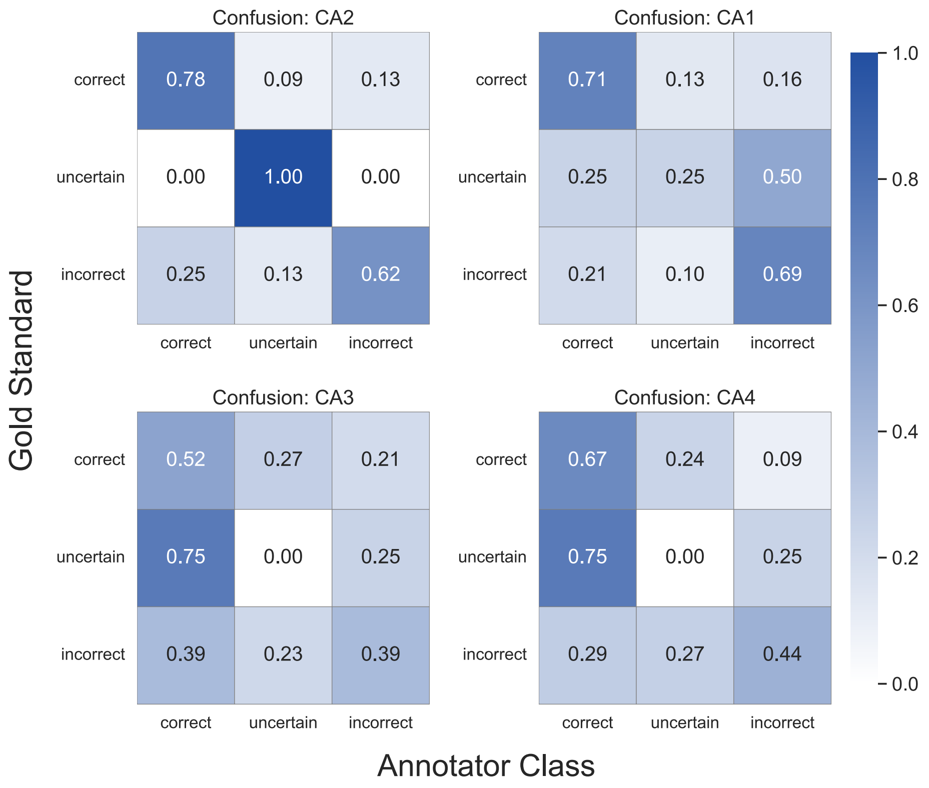


Figure S1: **Confusion matrix between** `gs` **and calibrated annotators for each classification categories.** The four confusion matrices are computed against the expert (`gs`) as ground truth. Diagonals vary by annotator and class; e.g., for `CA1`, items that are *correct* under the expert are labelled *correct* $\sim$71% of the time, and *incorrect* items are labelled *incorrect* $\sim$0.69 of the time. For `CA2`, the *uncertain* row is perfectly matched (1.00) on the diagonal in this sample, with lower diagonals for *correct* ($\sim$0.78) and *incorrect* ($\sim$0.62). These matrices make clear that disagreements are not uniform across classes or raters.

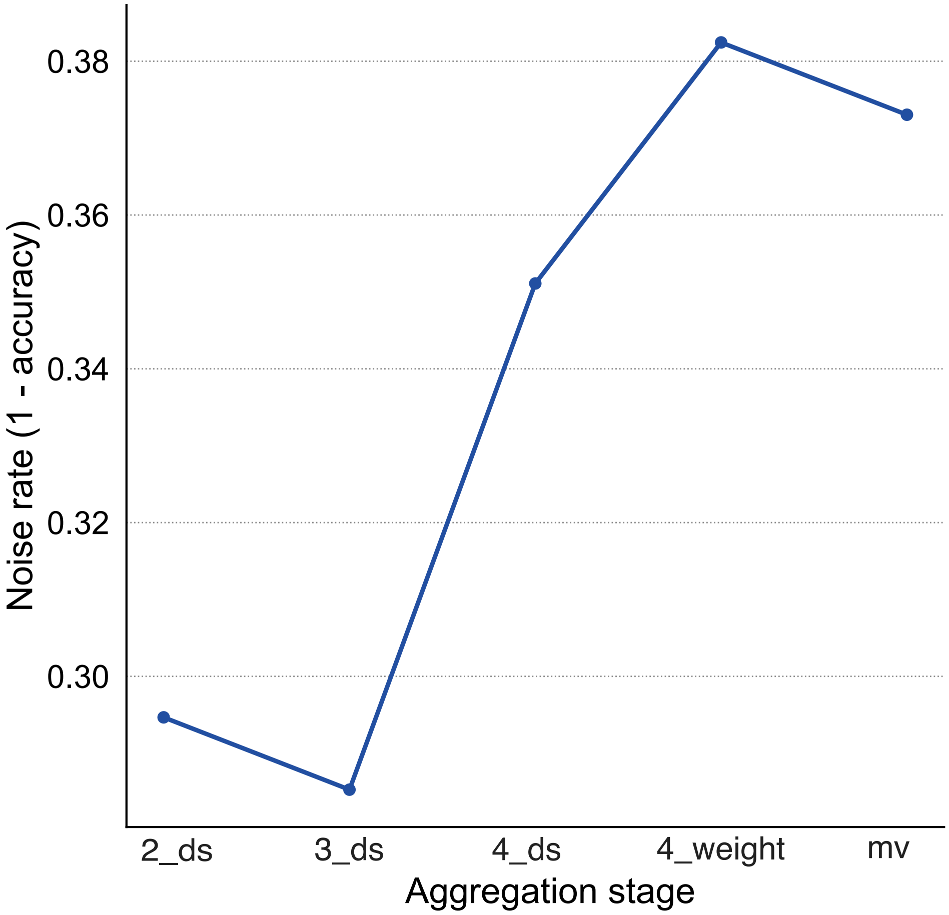


Figure S2: **Noise rate of various crowdsourced analysis methods:** Noise is defined as $1-$ weighted accuracy with respect to the gold standard. Dawid–Skene inferred from three annotators was closest to the gold standard, but increasing the number of annotators (**2_ds to 4_ds**) led to higher noise rates, further deviating from the gold standard. Weighted Dawid–Skene(**4_weight**) provides the furthest approximation to the gold standard than majority voting (**mv**) due to its emphasis on individual annotator judgments, resulting in the highest observed noise rate. Majority voting (**mv**) provided a reasonable approximation but still performed less than (**4_ds**)

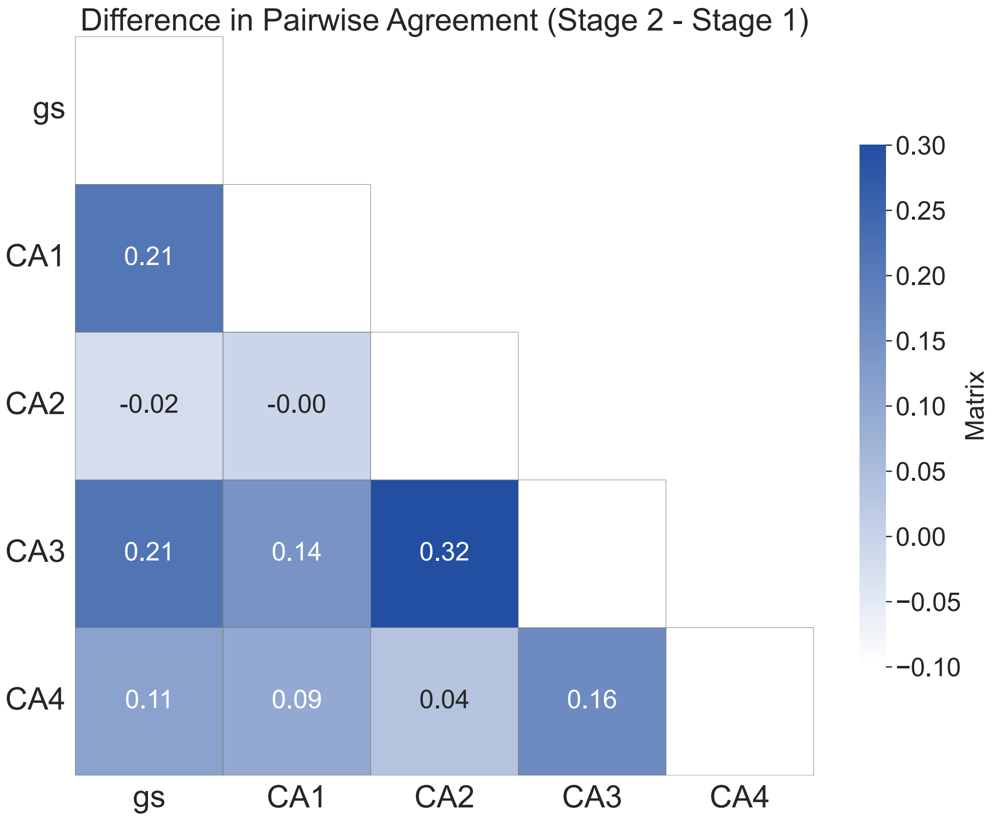


Figure S3: **Change in pairwise agreement (After Stage 2 - Before Stage 1)**. Heatmap of per-pair changes in agreement; positive values denote stronger post-calibration alignment. `CA3` shows the broadest improvement, increasing agreement with nearly all peers. `CA1` and `CA4` display smaller but mostly positive shifts. By contrast, `CA2` decreases in agreement with `gs` and shows limited gains with other raters, aside from a modest improvement with `CA3`. These patterns suggest that Stage 2 recalibration primarily benefited annotators who had adopted more conservative thresholds in Stage 1 (greater use of *uncertain*), bringing them closer to `gs`'s criteria, while highlighting one rater (`CA2`) whose decision boundary remained comparatively idiosyncratic.

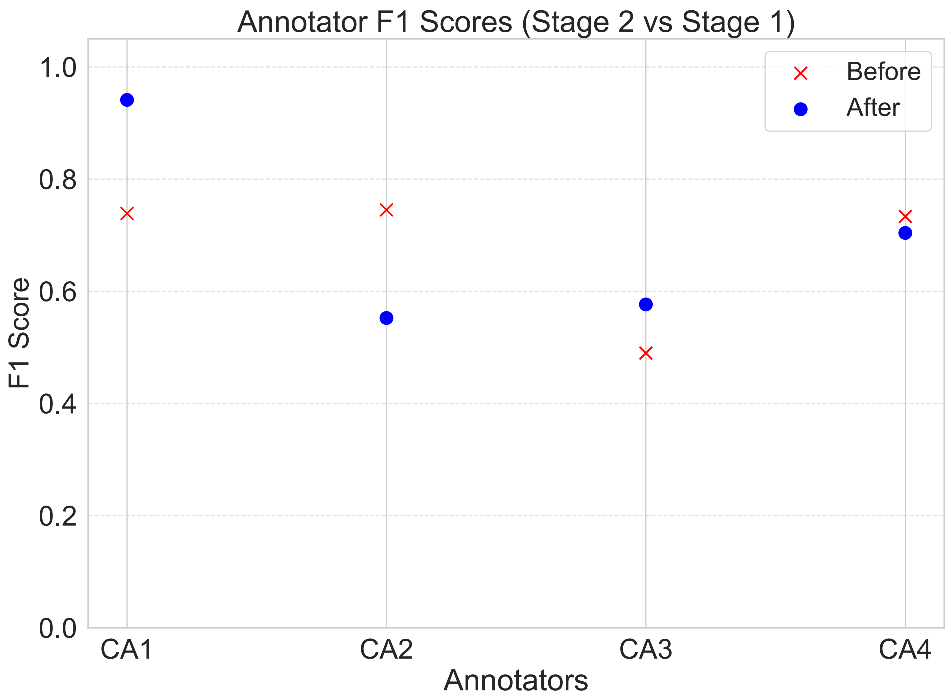


Figure S4: **Comparison of annotators' F1 scores before and after Stage 2 recalibration.** Only `CA1` and `CA3` showed improved F1 scores, with `acs` exhibiting the largest gain, indicating a greater ability to identify true positives after the pipeline. In contrast, both `syl` and `mr` experienced a decline in F1 scores, with `mr` showing the larger drop. This decline may reflect stricter annotation criteria, leading to reduced recall of positive synapse classifications.

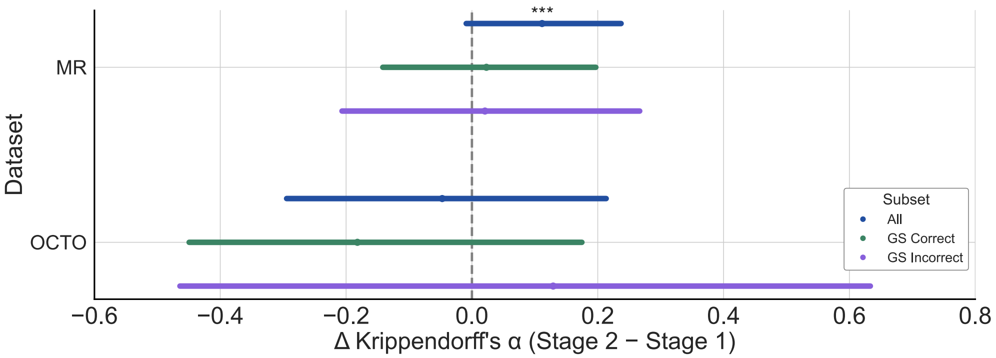


Figure S5: **Krippendorff's alpha differences before and after Stage 2 recalibration in MR and OCTO datasets.** A significant improvement in consensus was observed only in the MR dataset (***$p < 0.05$). Both datasets have subsets labeled as *incorrect* displaying moderate improvement in consensus than those labeled as *correct*. Compared to MR, the OCTO dataset exhibited a broader range of uncertainty and showed a decline in consensus.

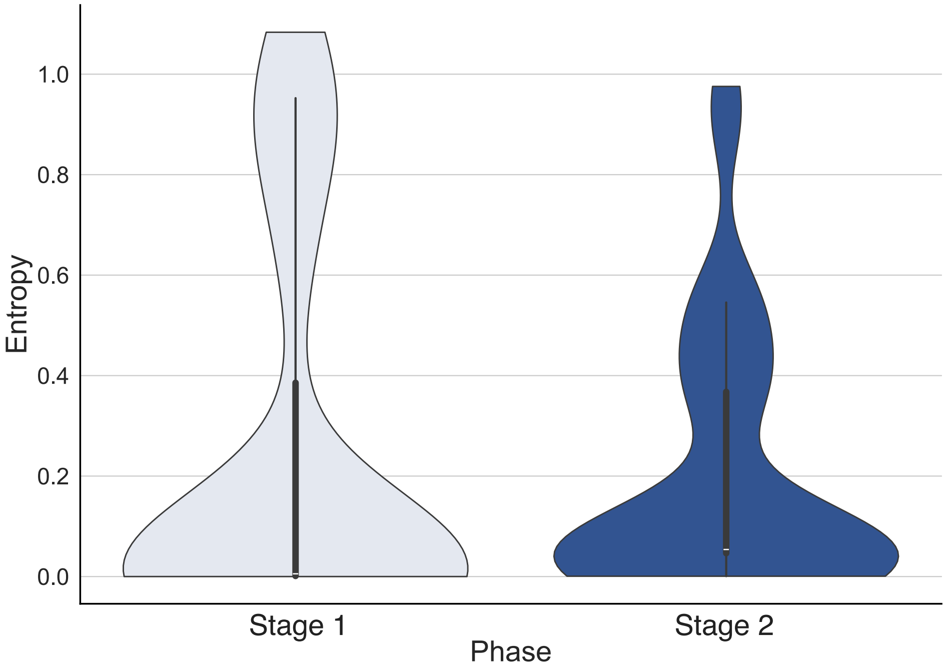


Figure S6: **Comparison of overall entropy distribution** Annotator uncertainty was higher before, with values concentrated near $\approx 1.0$, whereas in Stage 2, entropy was reduced and concentrated between 0 and 0.6, indicating greater agreement among CAs and increased confidence in synapse classification.

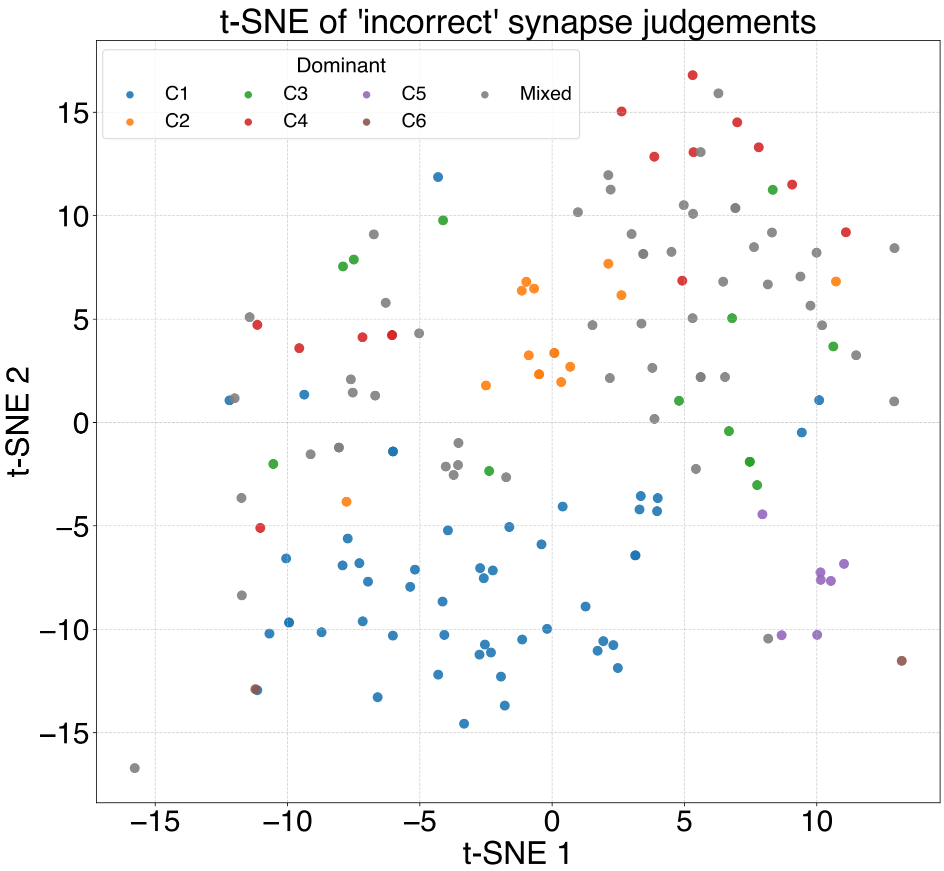


Figure S7: **t-SNE of features for synapses adjudicated *incorrect*.** Each point is a synapse judged *incorrect* in proofreading, embedded from the feature space into 2D with t-SNE. Points are coloured by the dominant rationale code: C1 = presynaptic T-bar absent/implausible; C2 = wrong postsynaptic partner; C3 = connector mispositioned relative to T-bar; C4 = postsynaptic node misassigned along $z$; C5 = self-synapse; C6 = imaging artefact; Mixed = no single dominant criterion. While classes overlap substantially, some locales (e.g., a central band enriched for C2) show partial grouping, suggesting that distinct feature signatures are associated with different error types. Axes denote the two t-SNE embedding dimensions.

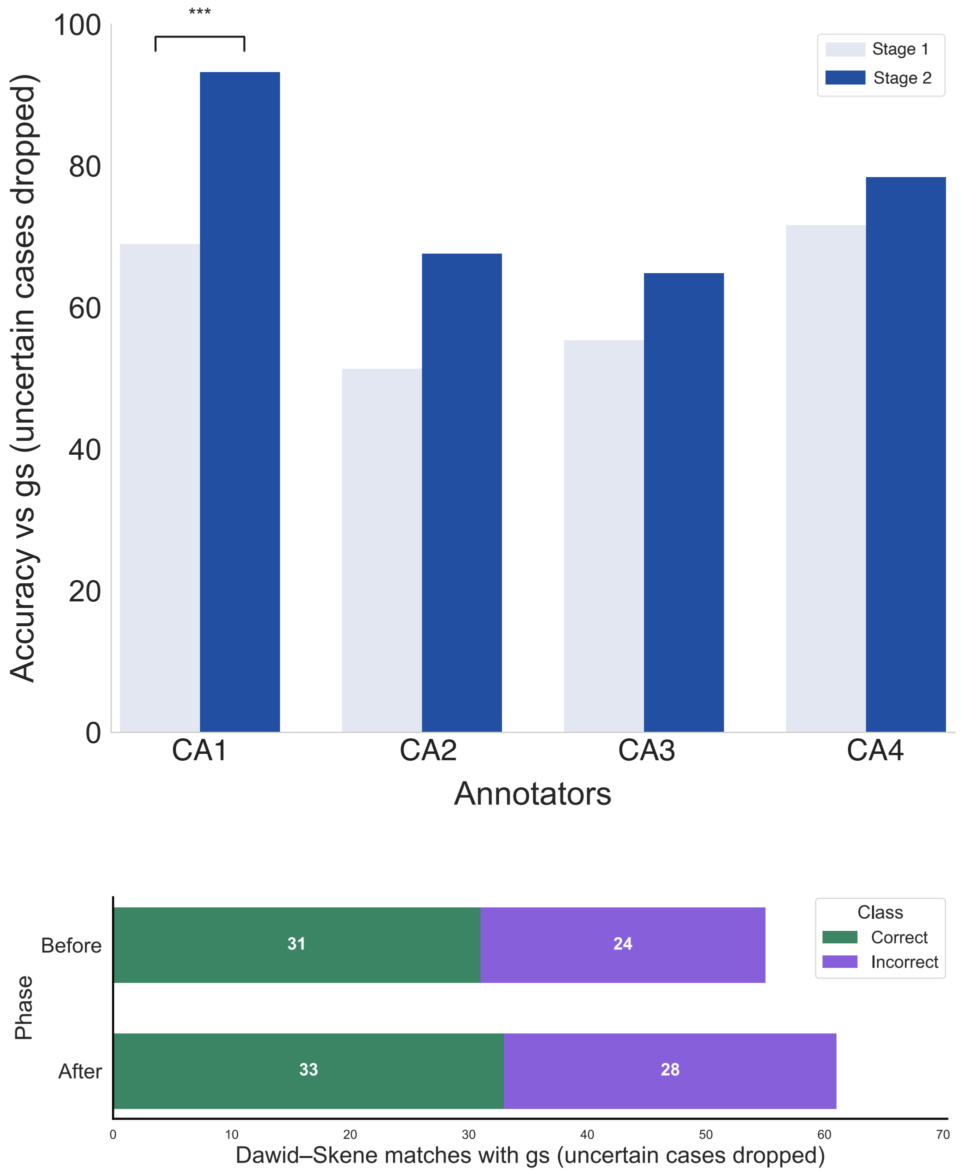


Figure S8: **Individual and DS consensus accuracy with *uncertain* cases dropped.** Across **73** paired cases without *uncertain*, no significant improvement was observed for most annotators (`CA2`: $p = 0.0501$; `CA3`: $p = 0.230$; `CA4`: $p = 0.267$), whereas only `CA1` retained a significant improvement ($p < 0.001$). DS analysis likewise indicated no significant change in overall crowd accuracy. Excluding the *uncertain* class encouraged adherence to the proofreading protocol, thereby improving individual and crowd-level alignment with gs
